# Comparative investigation into formycin A and pyrazofurin A biosynthesis reveals branch pathways for the construction of *C*-nucleoside scaffolds

**DOI:** 10.1101/728154

**Authors:** Meng Zhang, Peichao Zhang, Gudan Xu, Wenting Zhou, Yaojie Gao, Rong Gong, You-Sheng Cai, Hengjiang Cong, Zixin Deng, Neil P. J. Price, Xiangzhao Mao, Wenqing Chen

**Author notes:** These authors contributed equally to this paper (Meng Zhang, Peichao Zhang, Gudan Xu). For Correspondence: **Wenqing Chen**, School of Pharmaceutical Sciences, Wuhan University, Wuhan 430071, China. **E-mail**. For Co-correspondence: **Xiangzhao Mao**, College of Food Science and Engineering, Ocean University of China, Qingdao 266003, China. **E-mail**.

## Abstract

Formycin A (FOR-A) and pyrazofurin A (PRF-A) are purine-related *C*-nucleoside antibiotics, in which ribose and a pyrazole-derived base are linked by a *C*-glycosidic bond, however, the logic underlying the biosynthesis of these molecules has remained largely unexplored. Here, we report the discovery of the pathways for FOR-A and PRF-A biosynthesis from diverse actinobacteria, and demonstrate that their biosynthesis is initiated by a lysine *N*^6^-monooxygenase. Moreover, we show that the *forT* and *prfE* (individually related to FOR-A and PRF-A biosynthesis) mutants are correspondingly capable of accumulating the unexpected pyrazole-related intermediates, compound **11** and **9a**. We also decipher the enzymatic basis of ForT/PrfE for the *C*-glycosidic bond formation in FOR-A/PRF-A biosynthesis. To our knowledge, ForT/PrfE represents the first example of β-RFA-P (β-ribofuranosyl-aminobenzene 5’-phosphate) synthase-like enzymes governing *C*-nucleoside scaffold construction in natural product biosynthesis. These data establish a foundation for combinatorial biosynthesis of related purine nucleoside antibiotics, and also open the way for target-directed genome mining of PRF-A/FOR-A related antibiotics.

**IMPORTANCE:** Formycin A (FOR-A) and pyrazofurin A (PRF-A) are well known for their unusual chemical structures and remarkable biological activities. Actually, deciphering FOR-A/PRF-A biosynthesis will not only expand biochemical repertoire for novel enzymatic reactions, but also permit the target-oriented genome mining of FOR-A/PRF-A related *C*-nucleoside antibiotics.

## INTRODUCTION

Naturally-occurring *C*-nucleosides, e.g. formycin A (FOR-A) and pyrazofurin A (PRF-A) (Fig. 1) of microbial origin, are widely-distributed biological molecules in which the base and ribosyl moieties are linked *via* a structurally conserved *C*-glycosidic bond (1, 2). Pseudouridine, the first discovered *C*-nucleoside, is the most prevalent of the over one hundred different modified nucleosides found in eukaryotic RNA (3). Moreover, a pseudouridine metabolic pathway from *E. coli* has been previously reported and well-characterized (4). Also an important part of this group is the potent *C*-nucleoside antibiotics, which include the purine-related FOR-A and PRF-A (2), and the pyrimidine-derived showdomycin (5) and pseudouridimycin (Fig. 1) (6). The pathway for showdomycin biosynthesis was discovered relying on the enzyme pair AlnA (*C*-glycosyltransferase) and AlnB (phosphatase) as a probe, which catalyzes the formation of the *C*-ribosylated aromatic polyketide alnumycin C in a two-step process (5). Recently, Sosio *et al*, identified the pseudouridimycin biosynthetic gene cluster by searching for the gene (contained in a gene cluster) encoding pseudouridine synthase (6).

**Figure 1.**
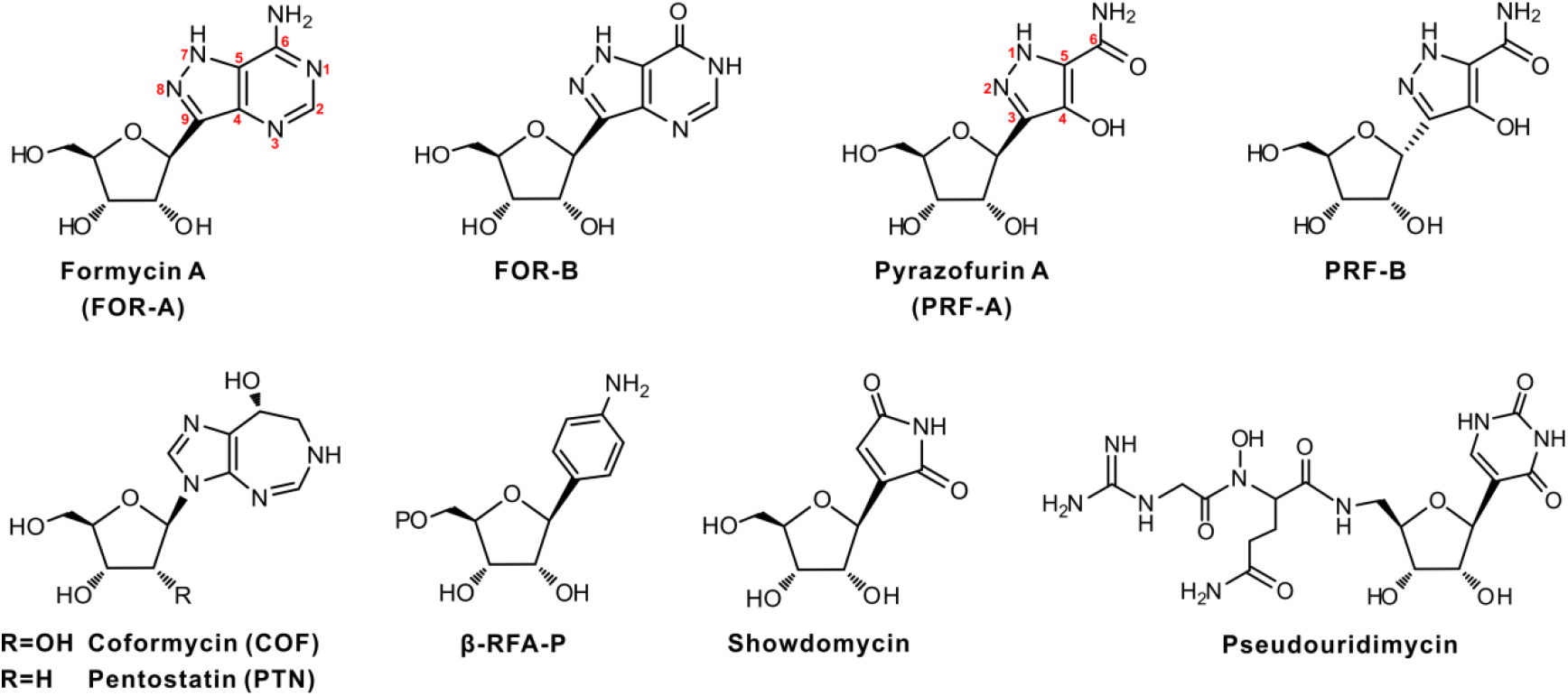
Chemical structures of FOR-A, PRF-A, and related compounds. β-RFA-P, β-ribofuranosyl-aminobenzene 5’-phosphate.

FOR-A was originally isolated from the broth of *Nocardia interforma* in the process of screening for antitumor compounds (7), and later, this antibiotic, accompanied by FOR-B (a deaminated product of FOR-A) (Fig. 1), was discovered to be produced by *Streptomyces lavendulae* (8). FOR-A and FOR-B (particularly the latter) show bioactivities against *Xanthomonas oryzae*, a pathogen incurring a rice plant disease, and the influenza A virus, while only FOR-A shows antitumor and antiviral activities (9). Subsequently, FOR-A and coformycin (COF, an adenosine deaminase inhibitor harboring a 1,3-diazepine ring) were found to be concomitantly produced by *Nocardia interforma* ATCC 21072 and *Streptomyces kaniharaensis* SF-557 (ATCC 21070) (Fig. 1) (10). This intriguing co-biosynthetic phenomenon was also documented for other purine nucleoside antibiotics pairs, including pentostatin (PTN) and arabinofuranosyl adenine (Ara-A) (produced by *Streptomyces antibioticus* NRRL 3238) (11), 2’-Cl PTN and 2’-amino-2’-deoxyadenosine (produced by *Actinomadura* sp. ATCC 39365) (12), and fungus-produced PTN and cordycepin (13), which all employ an unusual but general protector-protégé strategy, i.e. PTN may protect Ara-A (taking this antibiotic-pair as example) from deamination by the housekeeping adenosine deaminase (11–13).

PRF-A, produced by *Streptomyces candidus* NRRL 3601, possesses prominent antiviral and antitumor activities, but interestingly, PRF-B (Fig. 1), the α-anomer of PRF-A co-produced by this strain at a less amount, shows little bioactivity (14). Previous elegant studies indicated that PRF-A blocks *de novo* biosynthesis of pyrimidine and purine by independently targeting orotidylate decarboxylase (15) and 5-aminoimidazole-4-carboxamide ribonucleotide (AICAR) formyltransferase (16).

Earlier isotope feeding experiments have demonstrated that glutamate and ribose play key roles in *C*-glycosidic bond formation of FOR-A, and indicated that the C-3 to C-6 unit of PRF-A (C-9, C-4, C-5, and C-6 of FOR-A) is derived from C-4 to C-1 of glutamate and/or α-ketoglutarate (17). Moreover, Ochi *et al*, confirmed that N-3, N-7, and N-8 of FOR-A originate from the ε-amino nitrogen of lysine (18). More recently, Wang *et al*, identified the FOR-A biosynthetic gene cluster on the basis of previous studies (19), and deciphered the tailoring steps to FOR-A biosynthesis, by which they have established that FOR-A biosynthesis shares a crosstalk with the *de novo* purine pathway (19). Although a pathway for FOR-A biosynthesis was tentatively proposed, several important aspects have still remained unveiled.

In the present work, we report the identification and comparative analysis of the FOR-A/PRF-A biosynthetic gene clusters from diverse actinobacteria, and further show that the biosynthesis of FOR-A/PRF-A is initiated by a lysine *N*^6^-monooxygenase. We also demonstrate that a β-RFA-P synthase-like enzyme governs the construction of *C*-glycosidic bond of FOR-A/PRF-A associated with a cryptic decarboxylation.

## RESULTS

### Identification of the FOR-A gene clusters from diverse actinobacteria

FOR-A and COF were formerly reported to be co-produced by the actinobacteria, *Nocardia interforma* ATCC 21072 and *S. kaniharaensis* ATCC 21070 (7, 10), implicating that the genes for their biosyntheses, as those of other purine nucleoside antibiotic pairs, should also constitute a gene cluster (11, 12), Additionally, COF and PTN, featuring significant structural resemblance, were recently revealed to share highly-similar biosynthetic pathways (20).

To identify the target gene clusters for FOR-A biosynthesis, we sequenced the genomes of both strains using the Illumina method, which rendered ca. 9.3-Mb *(Nocardia interforma* ATCC 21072) and ca. 8.6-Mb (*S. kaniharaensis* ATCC 21070) of non-redundant bases after assembly of clean reads. We then utilize PenB (short-chain dehydrogenase) and PenC (SAICAR synthetase), the key enzymes for PTN biosynthesis, as probes (11), and the target gene clusters encoding the enzymes denoted as FocJ/CofA (66%/49% identity to PenB), and FocI1/CofB (69%/62% identity to PenC) (Fig. 2A, Table S1), respectively, were detected by genome mining of *Nocardia interforma* ATCC 21072 and *S. kaniharaensis* ATCC 21070. To correlate the *foc* gene cluster to FOR-A biosynthesis, we deleted a 1.8-kb region (containing *focH, focI1*, and *focJ*) (Fig. S1A). After confirmation by PCR analysis (Fig. S1B), the resultant mutant was fermented for metabolite analysis, and the HPLC and LC-HRMS analyses indicated that the production of FOR-A and COF was abolished in the PC1 mutant (Fig. 2B, C, Fig. S1C), confirming the identity of the *foc* gene cluster for the biosynthesis of the COF-FOR-A pair.

**Figure 2.**
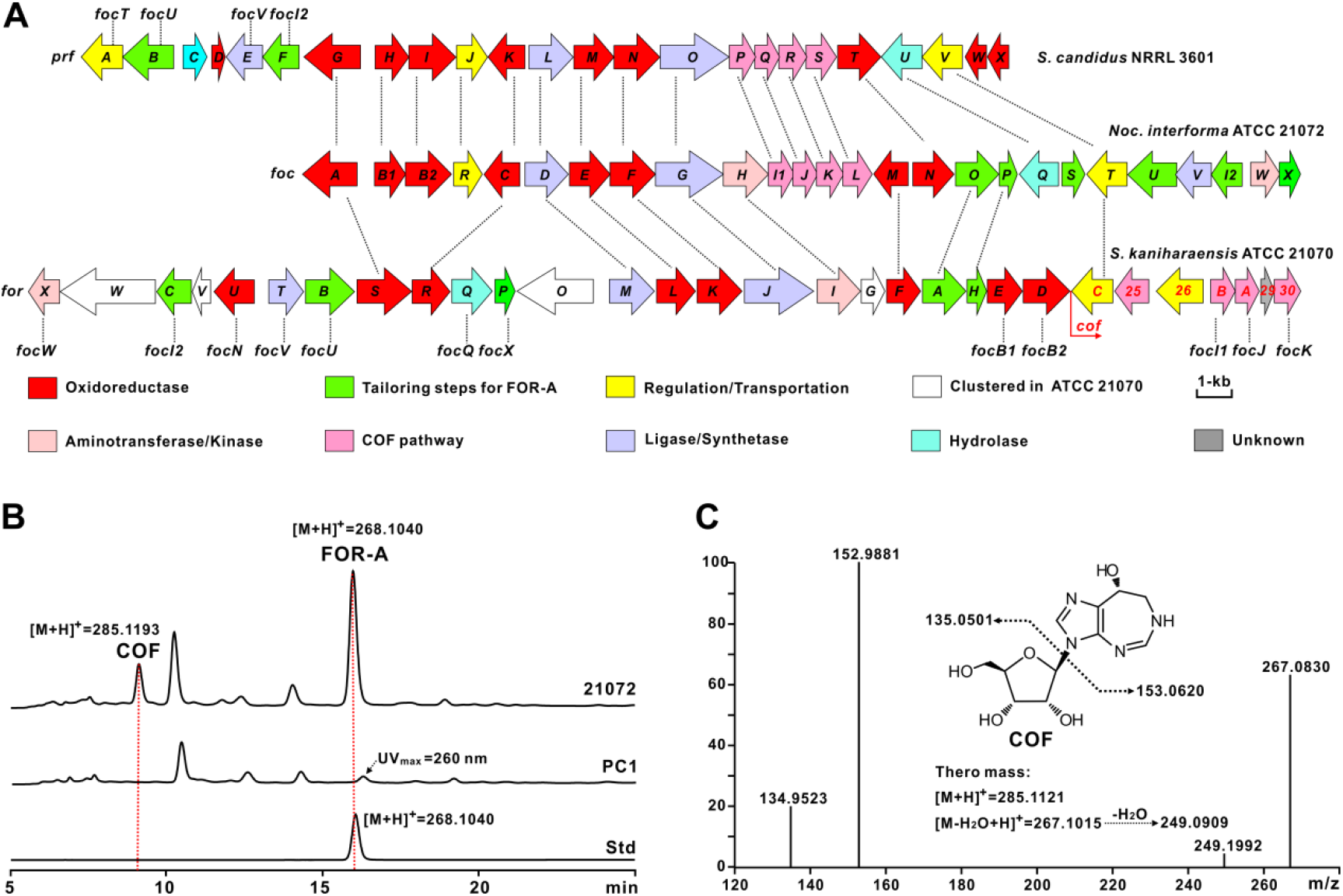
Genetic organization and verification of the PRF-A/FOR-A and COF (related) gene cluster. (A) Genetic organizations of the PRF-A/FOR-A and COF (related) gene cluster. The PRF-A gene cluster (*prf*) is from *S. candidus* NRRL 3601, and the FOR-A and COF gene cluster (*foc*) is from *Nocardia interforma* ATCC 21072; The genes responsible for FOR-A and COF biosynthesis (*for* and *cof*) in *S. kaniharaensis* ATCC 21070 are also designated as *cof1-30* (from left to right, Accession no. GenBank KY705052); The genes for COF biosynthesis in ATCC 21070 are marked by red, and the designation *cof* was used for the consistency with the study by wang *et al* (but *cof25, cof26, cof29, and cof30* are missed or not denoted in this work);^19^ In addition, the *foc* genes at the top correspond to the *prf* genes, by which the encoded enzymes are individually homologous; *Noc. interforma* ATCC 21072, *Nocardia interforma* ATCC 21072. The genes coding for homologous enzymes in these three gene clusters are correspondingly marked by dotted line. (B) Verification of the gene cluster (*foc*) for the biosynthesis of FOR-A and COF in *Nocardia interforma* ATCC 21072. 21072, the metabolites of *Nocardia interforma* ATCC 21072; PC1, the metabolites of *Nocardia interforma* PC1, in which 1.8-kb region of the *foc* gene cluster was deleted; Std, the authentic standard of FOR-A.

Further bioinformatic analysis led to the discovery of several potential FOR-A gene clusters with organizational diversities from other actinobacteria strains, including *Streptomyces resistomycificus* NRRL 2290 and *Salinispora arenicola* CNS-205 (Fig. S2A). We then fermented both of the strains for metabolites analysis, and the HPLC and LC-MS results indicated that *S. resistomycificus* NRRL 2290, but not *Salinispora arenicola* CNS-205, is capable of synthesizing FOR-A under the chosen fermentation conditions (Fig. S2B-D). These data suggested that the FOR-A gene clusters highlighting organizational diversities are widely-distributed among actinobacteria, but only partial strains are conferred with the capability of FOR-A production.

### Discovery of the PRF-A gene cluster from *S. candidus* NRRL 3601

The PRF-A molecule is structurally similar to FOR-A, implicating that both of them should employ identical or similar enzymatic logic for the construction of the pyrazole ring. To search for the candidate gene cluster for PRF-A biosynthesis, we sequenced the genome of *S. candidus* NRRL 3601, generating ca. 8.3-Mb non-redundant bases after assembly of orderly data, and then we used FocG (methionine-tRNA ligase) as an enzyme probe to perform BlastP analysis. This approach leads to the location of a candidate gene cluster from the genome of *S. candidus* NRRL 3601, encoding PrfO (86%/58% identities to FocG/ForJ), and neighboring genes PrfN (L-lysine *N*^6^-monooxygenase, 84%/62% identities to FocF/ForK) and PrfP (SAICAR synthetase, 82%/60% identities to FocI1/CofB). This strongly suggested that the target gene cluster is involved in PRF-A biosynthesis (Fig. 2A, Tables 1, S1).

**Table 1.** Deduced functions of the open reading frames in the *prf* gene cluster

| Protein | aa | Protein Function | Homolog, Origin | Identity, similarity (%) | Accession no. |
| --- | --- | --- | --- | --- | --- |
| PrfA | 396 | MFS transporter | NbrT6, *Nocardia terpenica* | 72, 84 | AJO72759 |
| PrfB | 476 | adenylosuccinate lyase | Orf67, *Nocardia terpenica* | 81, 89 | AJO72760 |
| PrfC | 224 | HAD family hydrolase | CLW14_2649, *Streptomyces* sp. 75 | 56, 66 | REE32254 |
| PrfD | 118 | cupin domain containing protein | CRH09_08295, *Nocardia terpenica* | 89, 94 | ATL66197 |
| PrfE* | 334 | β-RFA-P synthase-like enzyme | Orf61, *Nocardia terpenica* | 81, 87 | AJO72754 |
| PrfF | 343 | SAICAR synthetase | Orf62, *Nocardia terpenica* | 76, 84 | AJO72755 |
| PrfG | 540 | fumarate reductase/succinate dehydrogenase | Orf63, *Nocardia terpenica* | 77, 83 | AJO72756 |
| PrfH | 329 | putative monooxygenase | Sare_0593, *Salinispora arenicola* CNS-205 | 55, 64 | ABV96521 |
| PrfI | 444 | putative monooxygenase | Orf75, *Nocardia terpenica* | 79, 86 | AJO72768 |
| PrfJ | 288 | transcriptional regulator | NbrR10, *Nocardia terpenica* | 59, 70 | AJO72769 |
| PrfK | 351 | glycine/D-amino acid oxidase | EV381_1130, *Actinopolyspora* sp. DSM 45956 | 70, 79 | RZU68182 |
| PrfL | 413 | phosphoribosylglycinamide synthetase | Orf72, *Nocardia terpenica* | 77, 86 | AJO72765 |
| PrfM | 390 | saccharopine dehydrogenase | Orf71, *Nocardia terpenica* | 66, 77 | AJO72764 |
| PrfN* | 437 | L-lysine $N^6$-monooxygenase | Spb38, *Streptomyces* sp. SoC090715LN-17 | 42, 58 | BAW27702 |
| PrfO | 668 | methionine-tRNA ligase | Spb40, *Streptomyces* sp. SoC090715LN-17 | 33, 47 | BAW27704 |
| PrfP | 250 | SAICAR synthetase | PenC, *S. antibioticus* | 69, 79 | AKA87338 |
| PrfQ | 233 | short chain dehydrognase | PenB, *S. antibioticus* | 66, 79 | AKA87339 |
| PrfR | 260 | hydrolase | DI270_030585, *Microbispora triticiradicis* | 39, 54 | RGA01201 |
| PrfS | 288 | ATP phosphoribosyltransferase | PenA, *S. antibioticus* | 49, 64 | AKA87340 |
| PrfT | 402 | phthalate 4,5-dioxygenase | Orf68, *Nocardia terpenica* | 79, 89 | AJO72761 |
| PrfU | 387 | amidohydrolase | Orf65, *Nocardia terpenica* | 77, 84 | AJO72758 |
| PrfV | 404 | MFS transporter | D5S19_24520, *Amycolatopsis panacis* | 85, 92 | RJQ80812 |
| PrfW | 156 | flavin reductase | C8D87_103611, *Lechevalieria atacamensis* | 72, 79 | RAS67272 |
| PrfX | 169 | FMN reductase | C8D88_10484, *Lechevalieria deserti* | 71, 82 | PWK86923 |

\*Function confirmed *in vitro* in this study.

### Comparative analysis of the gene clusters for FOR-A and PRF-A biosynthesis

Comparative analysis of the FOR-A gene clusters revealed that they indicate apparent diversities in genetic organization as well as in the total number of genes. The FOR-A gene cluster (30 genes) from *S. kaniharaensis* ATCC 21070 spans a 37.0-kb continuous chromosomal region, while the counterpart *foc* gene cluster (26 genes) from *Nocardia interforma* ATCC 21072 is ca. 29.4-kb in size (Fig. 2A, Tables 1, S1). As anticipated, *in silico* analysis indicated that the genes for FOR-A and COF biosynthesis are linked together as reported by a recent study (19). Moreover, there are 7 additional genes (*forW, forV, forO, forG, cof25, cof26*, and *cof29*) in *S. kaniharaensis* ATCC 21070 (Table S1), and it is proposed to execute some alternative functions during FOR-A and COF biosynthesis in this strain.

The *prf* gene cluster, as shown by bioinformatic analysis, is composed of 24 genes and approximately occupies a continuous 27.1-kb chromosomal region of *S. candidus* NRRL 3601. The homologous *prf* and *foc* gene clusters contain some specialized genes, respectively (Fig. 2A). Of these, the genes *prfC* (encoding HAD family hydrolase), *prfD* (encoding a cupin domain containing protein), *prfW* (encoding flavin reductase), and *prfX* (encoding FMN reductase) are only present in *prf* gene cluster (Table 1). Further analysis indicates that the *foc* gene cluster contains the special genes *focP* (AICAR transformylase) and *focO* (adenylosuccinate synthetase) for the tailoring steps of FOR-A biosynthesis (Table S1). More interestingly, the candidate genes *prfP, prfQ, and prfS* (corresponding to *focl1, focJ*, and *focL*), whose products are homologous to PenC, PenB, and PenA, respectively, from the PTN pathway, are also present in *prf* gene cluster. However, more surprisingly, we were unable to identify the potential COF-related compounds, either COF or PTN, from the culture broths of *S. candidus* NRRL 3601 under lab fermentation conditions. Moreover, two particular genes, *focM* (codes for a dehydrogenase) and *focH* (codes for an aminotransferase), are only existent in the FOR-A gene cluster, suggesting that both of them are involved in the dedicated enzymatic steps in FOR-A biosynthesis (Table S1).

### Biosynthesis of FOR-A and PRF-A is initiated by the lysine *N*^6^-monooxygenase ForK/PrfN

FOR-A and PRF-A feature a distinctive hydrazine moiety (*N-N* bond) (Fig. 1), which is also present in s56-p1, a dipeptide natural product produced by *Streptomyces* sp. SoC090715LN-17 (21). A recent study has revealed that the *N-N* bond construction is confirmatively governed by a cascade of two enzymes, Spb38 (lysine *N*^6^-monooxygenase) and Spb40 (methionine-tRNA ligase-like protein) (Table 1, Fig. 3A) (21). Bioinformatic analysis of the FOR-A and PRF-A gene cluster rendered the identification of two candidates ForK/PrfN (deduced lysine *N*^6^-monooxygenase) and ForJ/PrfO (methionine-tRNA ligase-like protein) (Table S1, Fig. S3). To determine if ForK/PrfN fulfils the potential role of lysine *N*^6^-monooxygenase for the initiation of FOR-A/PRF-A biosynthesis, we overexpressed and purified the protein in *E. coli* (Fig. S4A). As expected, the ForK/PrfN protein displays color of light yellow (Fig. S4B), a typical characteristic of flavoprotein (Fig. S5). We then tested their enzymatic activity *in vitro* with L-lysine as substrate and NADPH as cofactor. As anticipated, LC-MS analysis indicated that the ForK/PrfN reaction mixtures are capable of generating a distinctive [M+H]^+^ ion of *N*^6^-OH lysine at *m/z* 163.1088/163.1079, with three major fragment ions at *m/z* 99.8577/99.9317, 129.9084/129.9299, and 144.8742/144.9361 (Fig. 3B, Fig. S6A-B), which are completely consistent with the fragmentation pattern of the positive control (the NbtG reaction) (Fig. S6C-D, Table S4) (22). However, the reaction (without enzyme added as negative control) mixtures, as detected by LC-MS, could not produce the expected MS peaks (Fig. 3B). Moreover, we tested the cofactor specificity for the two enzymes, and the LC-MS results indicated that the reaction mixtures with NADH as cofactor is also able to give the anticipated [M+H]^+^ ion of *N*^6^-OH lysine, confirming that NADH could also support the activity of PrfN/ForK (Fig. S7). Taken together, these enzymatic data established that PrfN/ForK is undoubtedly a lysine *N*^6^-monooxygenase for the initiation of PRF-A/FOR-A biosynthesis.

**Figure 3.**
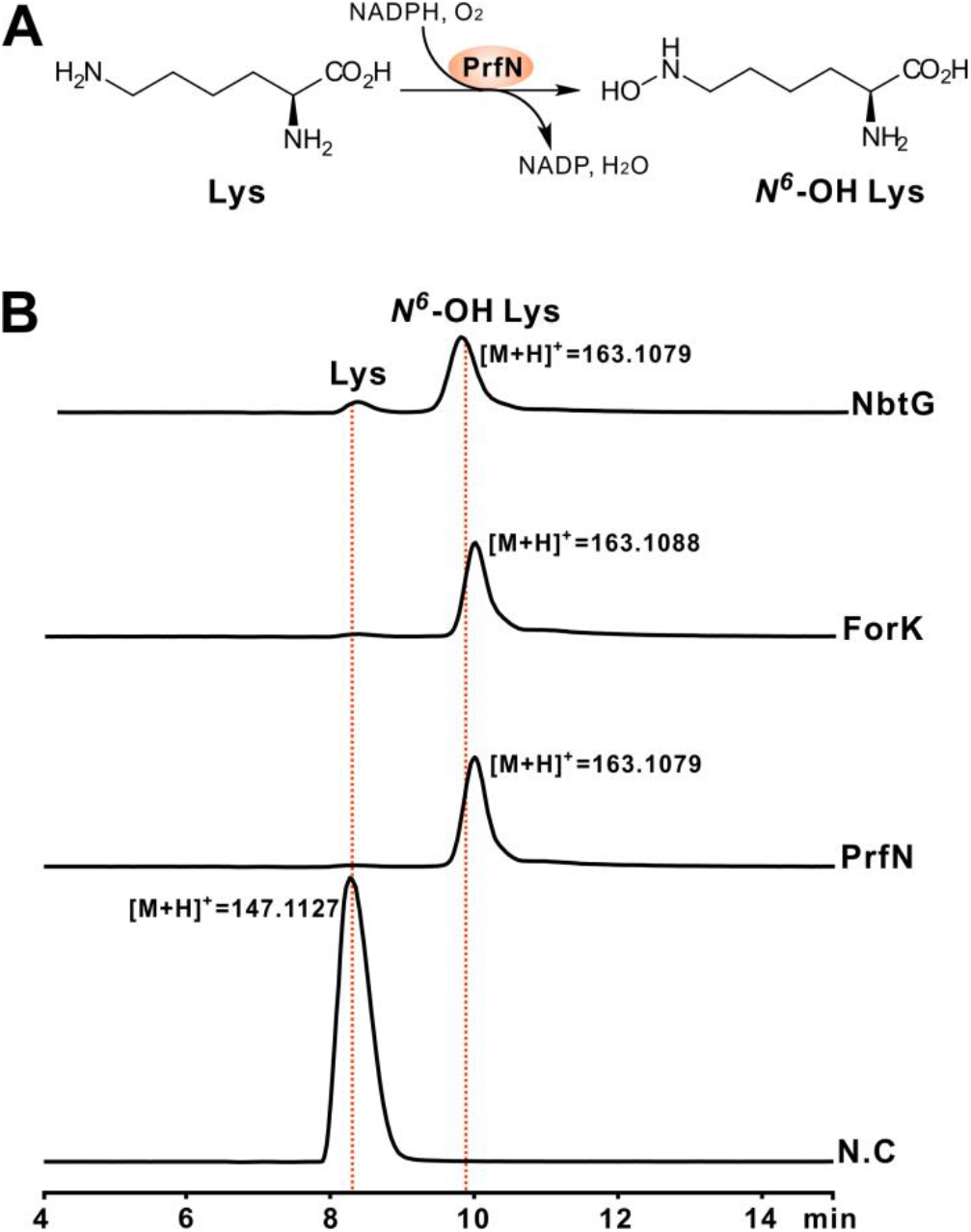
Biochemical characterization of ForK/PrfN as a lysine *N*^6^-monooxygenase. (A) Scheme of the ForK/PrfN-catalyzed reaction. Lys, Lysine; (B) LC-MS analysis of the ForK/PrfN reaction using Lysine (Lys) as substrate and NADPH as cofactor. NbtG, the NbtG reaction as positive control, NbtG (Acession no. BAD55606) is from *Nocardia farcinica* IFM 10152; ForK, the ForK reaction; PrfN, the PrfN reaction; N.C, the reaction without enzyme added as negative control.

### The *ΔforT* and *ΔprfE* mutants accumulate the corresponding intermediates, compound 11 and 9a

FOR-A/PRF-A features an unusual *C*-nucleoside scaffold, but how nature constructs the *C-C* bond has long remained mysterious. *In silico* investigation of the FOR-A/PRF-A gene cluster revealed that *forT/prfE* encodes a deduced β-ribofuranosyl-aminobenzene 5’-phosphate synthase (Tables 1, S1). To investigate the *in vivo* functional role of *forT/prfE*, we in-frame deleted it using a CRISPR-Cas9 strategy (Fig. S8A-B, S9A-B) (23). As *S. kaniharaensis* ATCC 21070 proved more amenable to in-frame deletion manipulation than *Nocardia interforma* ATCC 21072, it was used in this study for the following genetic experiments. After confirmation by a combined PCR and sequencing analysis (Fig. S8B, Table S5), the mutant *ΔforT* was fermented for metabolites analysis. HPLC analysis indicated that the metabolites of the *ΔforT* mutant could not generate the distinctive FOR-A peak, while capable of yielding a new one (compound **11**) at RT = 15.8 min with UV_max_ = 308 nm (Fig. 4A, Fig. S8C-E). LC-MS analysis of compound **11** indicates that it shows an apparent [M+H]^+^ ion at *m/z* 172.0353 (Fig. 4A), which could produce a specific fragment ion at *m/z* 153.8110 (Fig. S8F). Likewise, the confirmed *ΔprfE* mutant (Fig. S9B, Table S5) lacks the capability of PRF-A and PRF-B production (Fig. S9C-D), but its metabolites could generate an obvious new peak (compound **9a**) at RT = 22.5 min with UV_max_ = 273 nm (Fig. 4B, Fig. S9E). LC-MS analysis of the compound **9a** peak indicates that it shows an apparent [M+H]^+^ ion at *m/z* 173.0193 as well as the main fragment at *m/z* 154.0294 (Figs. 4B, S9F).

**Figure 4.**
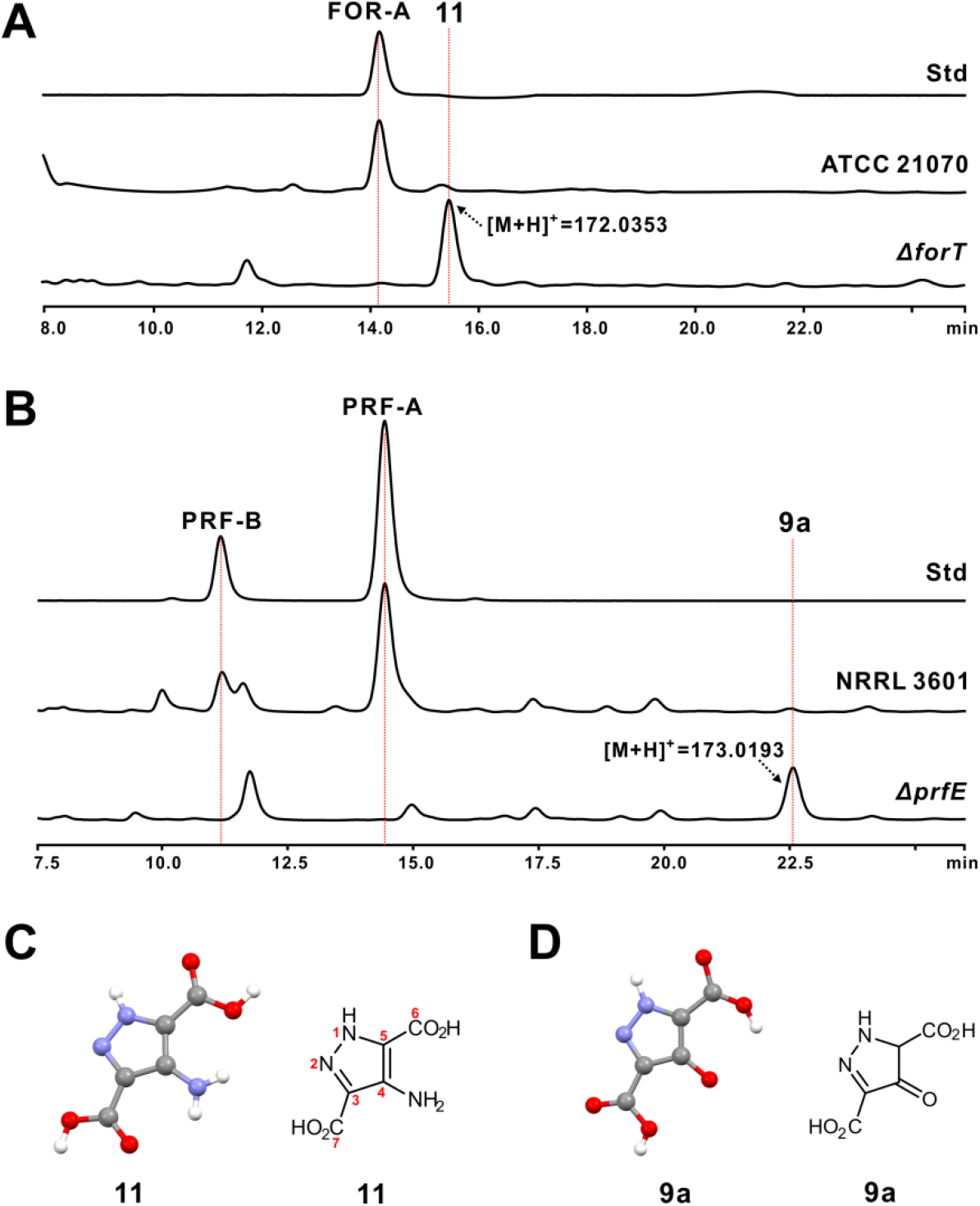
HPLC analysis of the metabolites produced by the *ΔforT* and *ΔprfE* mutants. (A) HPLC analysis of the metabolites produced by *S. kaniharaensis ΔforT* mutant. Std, the authentic standard of FOR-A; ATCC 21070, the metabolites produced by wild-type of *S. kaniharaensis* ATCC 21070; *ΔforT*, the metabolites of *ΔforT* mutant. (B) HPLC traces of the metabolites produced by *the S. candidus ΔprfE* mutant. Std, the authentic standard of PRF-A; NRRL 3601, the metabolites produced by the wild-type of *S. candidus* NRRL 3601; *ΔprfE*, the metabolites of *ΔprfE* mutant. (C) Crystal and chemical structures of the compound **11**. (D) Crystal and chemical structures of the compound **9a**. The numbering for the atoms of **9a** corresponds to those of **11**.

To determine the chemical structures of compounds **11** and **9a**, we initially purified them *via* HPLC preparation for NMR analysis. However, neither of them gave rise to any apparently detectable ^1^H signals for structure elucidation, and we were therefore prompted to optimize the conditions of compound **11/9a** crystallization for X-ray structure determination (Fig. 4C-D, Tables S6, S7). To our surprise, we found that the compound **11/9a** harbors a highly-conjugated five-membered structural system, in which two *N*-atoms (N1 and N2) are linked together to form an unusual *N-N* bond (Fig. 4CD, Tables S6, S7). It is exactly because of the peculiar structure of **11/9a** that directly results in the almost undetectable ^1^H signals during NMR analysis. Taken together, these combined genetic and crystal data suggested that *prfE/forT* is most likely the candidate gene for the *C-C* bond formation during PRF-A/FOR-A biosynthesis.

### PrfE/ForT is a β-RFA-P synthase-like enzyme for the *C*-glycosidic bond formation during PRF-A/FOR-A biosynthesis

Bioinformatic analysis indicated that PrfE/ForT (Tables 1, S1) exhibits high homology (81%/64%) to Orf61 (hypothetic (β-RFA-P synthase) from *Nocardia terpenica* (Fig. S10). (β-RFA-P synthase is responsible for the first committed step of methanopterin biosynthesis, catalyzing the conversion of phosphoribosyl pyrophosphate (PRPP) and p-aminobenzoate (pABA) to (β-RFA-P (Fig. 1) and CO_2_ (24). To obtain direct biochemical evidence that PrfE/ForT governs the *C*-glycosidic bond construction during PRF-A/FOR-A biosynthesis, we overexpressed and purified these proteins from *E. coli* to near homogeneity and assayed them *in vitro* using PRPP and compound **9a/11** as substrates (Fig. S11A-B). The HPLC traces indicated that the product of the PrfE reaction is capable of producing a new peak at RT = 6.5 min with a distinctive UV_max_ = 267 nm (Fig. S12A), which is absent from that of the negative control (the reaction without PrfE added) (Fig. 5A, C, D). Further LC-MS analysis indicated that the product of the PrfE reaction exhibits an obvious [M+H]^+^ ion at *m/z* 341,0393 as well as the main fragment ions at *m/z* 224.9183, 242.8766, 322.9725, and others (Figs. 5A, S12E, I). Moreover, the reaction without the divalent metal Mg^2+^ is not able to generate the characteristic peak of the product. We therefore purified the target compound for further ^1^H, ^13^C, and 2D NMR analysis (Fig. S13-S17, and Table S8), and its chemical structure, on the basis of the combined NMR and MS/MS analysis, was finally determined as compound **19**.

**Figure 5.**
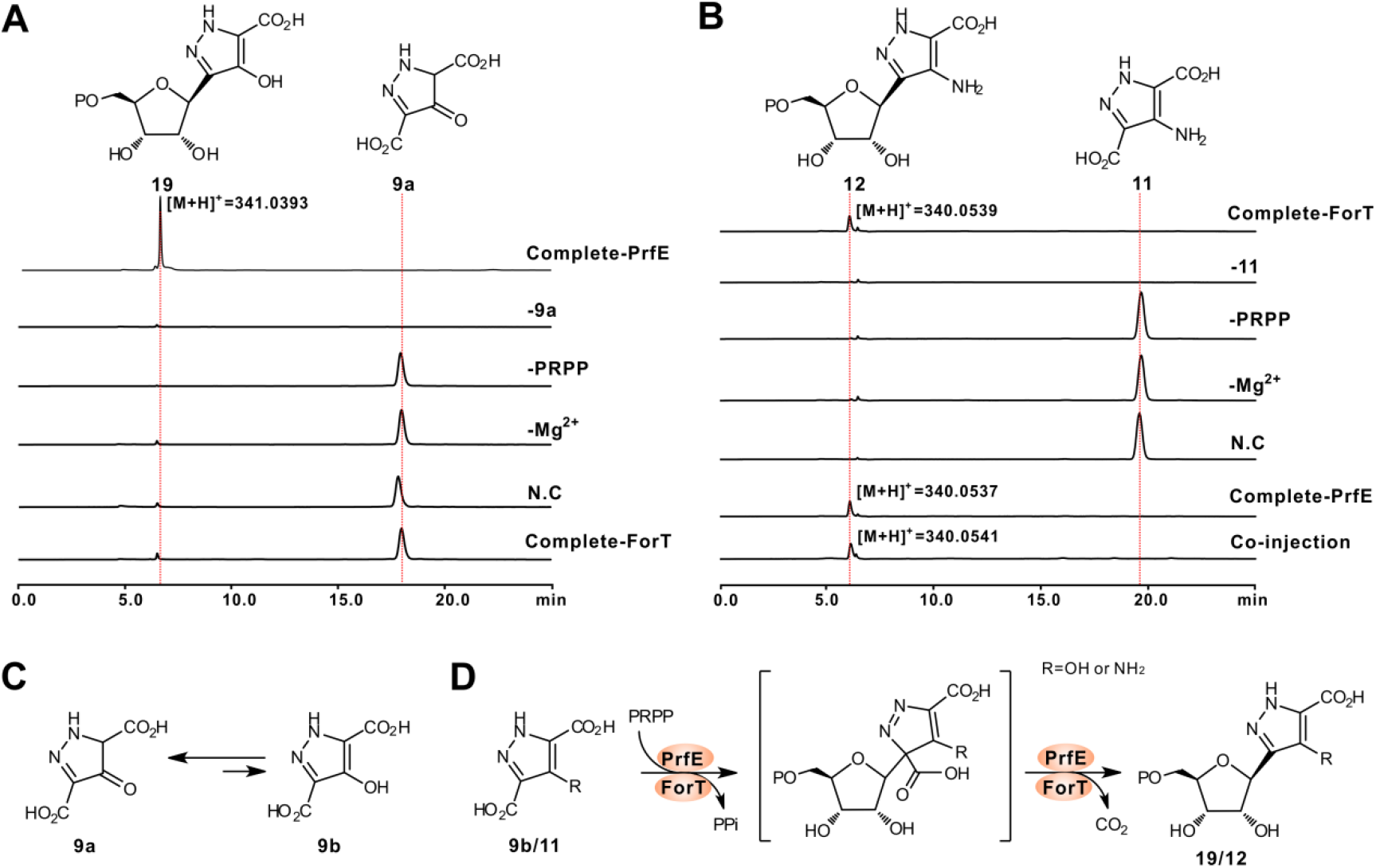
Biochemical characterization of PrfE and ForT as β-RFA-P synthase-like enzymes for the *C*-glycosdic bond construction. (A) HPLC traces of the PrfE-catalyzed reaction. Compete-PrfE, the complete reaction of PrfE using with **9a** and PRPP as substrates; −9a, the PrfE reaction without **9a** added; -PRPP, the PrfE reaction without **9a** added; -Mg^2+^, the PrfE reaction without Mg^2+^ added; N.C, the PrfE reaction lacking enzyme as the negative control; Compete-ForT, the complete reaction of ForT using with **9a** and PRPP as substrate. (B) HPLC traces of the ForT-catalyzed reaction. Compete-ForT, the complete reaction of ForT using **11** and PRPP as substrates; −11, the ForT reaction without **11** added; -PRPP, the ForT reaction without **11** added; -Mg^2+^, the ForT reaction without Mg^2+^ added; N.C, the ForT reaction without enzyme added as negative control; Compete-PrfE, the complete reaction of PrfE using with **11** and PRPP as substrates; Co-injection, co-injection of the ForT reaction and the PrfE reaction. (C) Scheme of the tautomerization reaction between **9a** and **9b**. (D) Proposed mechanism for the PrfE/ForT catalyzed reaction.

Similarly, the ForT reaction mixtures, as detected by HPLC analysis, are also capable of producing a characteristic peak (compound **12**) with a distinctive UV_max_ = 290 nm (Fig. 5B, S12B), which is distinctly absent from the reaction mixtures without ForT/Mg^2+^. Additional tandem MS/MS analysis showed that the target peak could produce an apparent [M+H]^+^ ion at *m/z* 340.0539 (Fig. 5B), with major fragments ions at *m/z* 205.9951, 223.8481, and 321.9420, completely consistent with the theoretical fragmentation pattern of compound **12** (Fig. S12F, J). Furthermore, we tested the substrate flexibility for PrfE, and the results indicated that this enzyme could also accept **11** to form **12** (Figs. 5B, S12C, D, G, H), but that conversely, ForT is not capable of recognizing **11** as substrate (Fig. 5A).

Subsequently, we examined the specificities of divalent metal ions for PrfE/ForT activity. Of the 6 divalent metal ions selected, Mn^2+^ and Co^2+^ are able to maintain the maximal activity for PrfE (100%), but it indicates either dramatically decreased activity in the presence of Ni^2+^/Zn^2+^, or negative activity with Ca^2+^/Cu^2+^ added instead (Fig. S12K). For the ForT reaction, Co^2+^ could also support its maximal activity, while Mn^2+^ (61%), Ni^2+^ (52%), Zn^2+^ (84%), or Ca^2+^/Cu^2+^ (0%) are just partially able or unable to take the role of Mg^2+^ to support the enzymatic activity of ForT (Fig. S12L). All of these biochemical data demonstrate that PrfE/ForT functions as an unusual β-RFA-P synthase-like enzyme for the *C*-glycosidic bond formation during PRF-A/FOR-A biosynthesis.

## DISCUSSION

Earlier metabolic labelling experiments determined that glutamate and lysine were the precursors for FOR-A and PRF-A biosynthesis, and the ribosyl moiety of both antibiotics was directly from the primary metabolism (17, 18). In the present study, this assignment is shown to be essentially correct. Insight into the biosynthetic gene cluster of PRF-A/FOR-A (only the PRF enzymes for common steps of both antibiotics are listed in Fig. 6) results in the identification of the enzymes for the construction of pyrazole ring system (Fig. 6A). PRF-A Biosynthesis is proposed to be initiated by PrfN catalyzing L-lysine to form *N*^6^-OH lysine, which, just as most lysine *N*^6^-monooxygenases, prefers NADPH as cofactor (22). Concomitantly, L-glutamate is dehydrogenated to form **1**, and then both intermediates are utilized by PrfO (methionine tRNA ligase) to construct the *N-N* bond in compound **3**, which is catalyzed by PrfK (amino acid oxidase) to afford **4** with leaving of compound **5**. Following on from this, we tentatively propose that compound **4** is hydroxylated by PrfH/I in a sequential manner at *C*-4 position to generate **6**, which would be successively dehydrated *via* either a spontaneous or an unknown enzymatic strategy to give rise to **7**. Further investigation of the PRF-A gene cluster results in the identification of a candidate enzyme PrfL (phosphoribosyl-glycinamide synthetase-like protein), which usually catalyzes the second step in *de novo* purine pathway (25). We therefore postulate that **7**, under the catalysis of PrfL, is subsequently converted to **8** for the accomplishment of the pyrazole ring construction. Thereafter, we propose that **8**, after oxidation to **9a**, will further undergo a series of sequential reactions to complete PRF-A biosynthesis (Fig. 6A).

**Figure 6.**
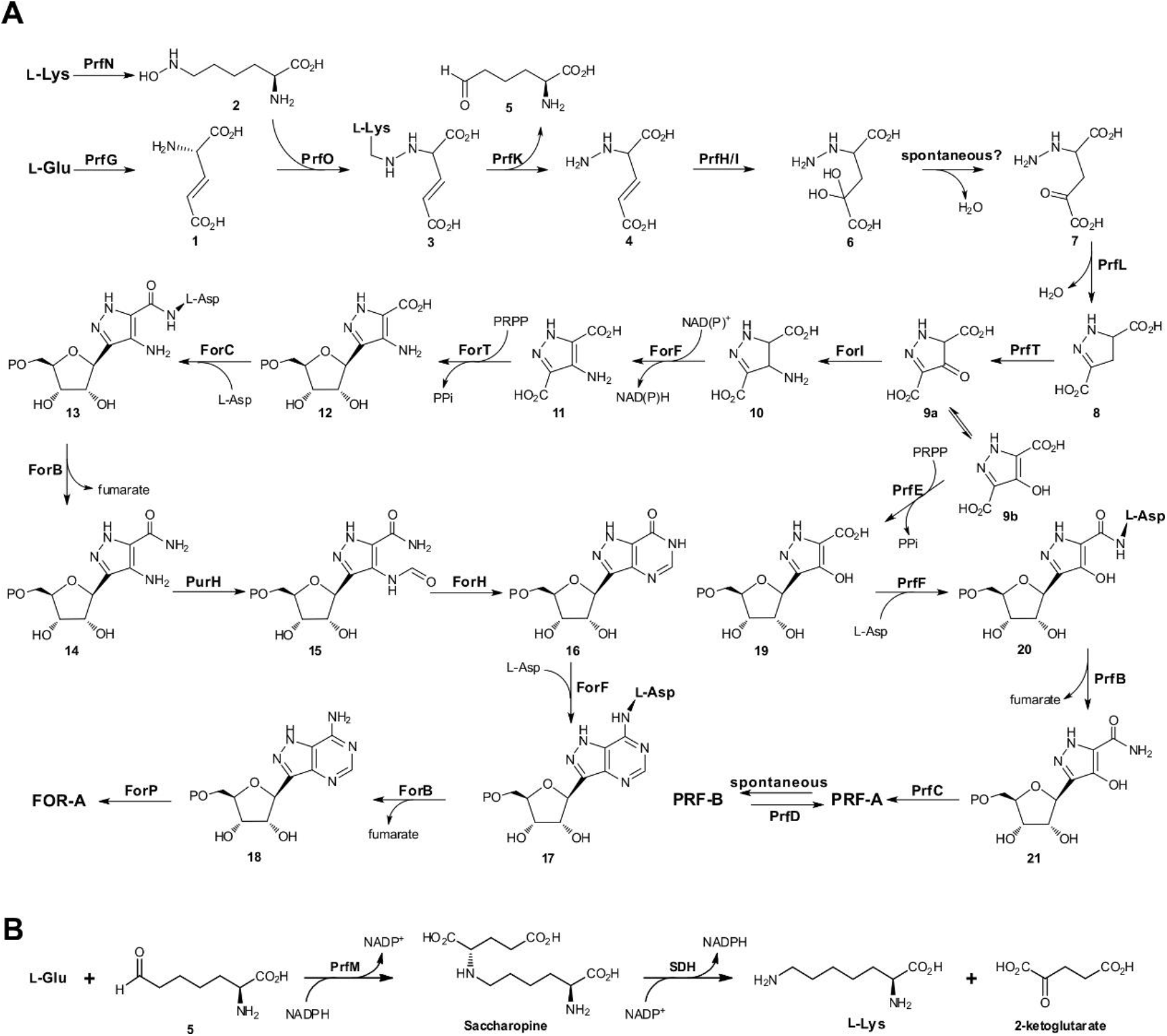
Proposed biosynthetic pathways to FOR-A and PRF-A. (A) Proposed pathways to FOR-A and PRF-A. We propose that they share the identical steps at the early stage prior to the construction of *C*-nucleoside scaffolds. Proposed salvage pathway for the regeneration of lysine during FOR-A/PRF-A biosynthesis. SDH, short chain dehydrogenase, which is from the primary metabolic pathway for lysine regeneration.

As for FOR-A biosynthesis, we deduce that **9a**, acting as a potential amino-group acceptor, will be modified by the aminotransferase ForI to **10**, which is then dehydrogenated to give the compound **11**. Actually, the aminotransferase (ForI) and dehydrogenase (ForF) are the particular enzymes absent from the PRF-A pathway, implying that they should perform these definite functions to accomplish **11** biosynthesis (Fig. 6A), and related biochemical studies in our laboratory are now ongoing. Once the *C*-glycosidic bond constructed, the later tailoring enzymatic steps are highly similar to those in the *de novo* purine pathway, which has already been illustrated in two recent studies (19, 26).

There are several other genes (*prfDMWX*) whose functional roles are still ambiguous. In particular, the product of *prfD* (cupin domain containing protein) is probably responsible for the conversion of PRF-B to PRF-A, although it is also possible to synthesize PRF-B from PRF-A *via* a spontaneous way. Hence, tentatively, PrfD is proposed to be an isomerase during PRF-A biosynthesis. PrfM is a potential saccharopine dehydrogenase, which normally plays a key role in lysine biosynthesis (27). Accordingly, we propose that PrfM could be responsible for a salvage pathway for lysine regeneration during PRF-A biosynthesis (Fig. 6B). For the enzymes, PrfW and PrfX, they are apparently homologous to flavin/FMN reductase, and could be involved in the reductive recycle of FMN/FAD cofactor in PRF-A biosynthesis.

*C*-Nucleoside antibiotics have arrested increasing interests for clinical uses in the past decades (2), but their biosynthetic logics have long been under-appreciated due to their difficult-to-access pathways. Actually, nature has developed diverse strategies for *C*-glycosidic bond construction during the biosynthesis of this group of antibiotics. Showdomycin biosynthesis uses a YeiN-like *C*-glycosyltransferase to build the *C-C* bond (5), while pseudouridimycin and malayamycin are deduced to exploit a tRNA (TruD-like) pseudouridylate synthase for *C*-glycosidic bond construction (6, 28). In the present study, FOR-A and PRF-A harness a totally different β-RFA-P synthase-like enzyme (ForT/PrfE) to catalyze the formation of *C*-glycosides. The usual role for β-RFA-P synthase has exclusively been in the biosynthesis of the modified folate methanopterin (Fig. S18A) (24). In this respect, ForT/PrfE represents the first example of this kind of enzymes responsible for the *C*-glycosidic bond formation in natural products biosynthesis. Hence, similar to β-RFA-P synthase (24), we predict that ForT/PrfE should employ a S_N_1-like enzymatic strategy associated with a cryptic decarboxylation for the construction of the *C*-glycosidic bond during PRF-A/FOR-A biosynthesis. Notably, the single β-RFA-P synthase-like enzyme (ForT/PrfE) could be used as a promising probe for the rational mining of related *C*-nucleoside antibiotics, and we have already discovered several potential PRF-A/FOR-A group antibiotic pathways from the currently-available reservoir of microbial genomes (Fig. S18B).

In summary, we report the pathways for FOR-A and PRF-A biosynthesis using a genomics-led approach, and have delineated the associated FOR-A gene-cluster diversity in actinobacteria. We have also demonstrated that biosynthesis of FOR-A and PRF-A is initiated by a lysine *N*^6^-monooxygenase, and show that PrfE/ForT adopts a unique strategy for the *C*-glycosidic bond formation in nucleoside antibiotics biosynthesis. We anticipate that the *C*-glycosyltransferases shown in this study may serve as an inspiration for future catalyst design to generate analogs of potential therapeutic value.

## MATERIALS AND METHODS

### General materials and methods

Strains, plasmids, and cosmids used in this study were described in Supplementary Table S1, and primers were listed in Supplementary Table S2. General methods employed in this work were according to the standard protocols of Green *et al* (29) or Kieser *et al* (30).

### Construction of the mutants, *Nocardia interforma* PC1, *S. kaniharaensis ΔforT*, and *S. candidus ΔprfE*

For the construction of the mutant *Nocardia interforma* PC1, a similar strategy by Wu et al (11), with Ni-lfdsaicar-LF/R (Left arm) and Ni-lfdsaicar-RF/R (Right arm) as primers and pOJ446 as starter vector, was also used in this study. For the construction of *S. kaniharaensis ΔforT* (taking this strain as example) mutant, the gRNA, after annealing, was cloned into the Xbal/Ncol-engineered site of pCRISPR-Cas9 (23) to generate pCas9-gRNA. Subsequently, the PCR products of left-arm, the right-arm, and the StuI-engineered pCas9-gRNA, amplified by KOD plus (TOYOBO), were recombined using the Multi-segment Mosaic Enzymes (Yeasen Biotech) to generate pCHW352 (Table S2), which was then introduced into *S. kaniharaensis via* conjugation. After confirmation, the conjugant was induced by thiostrepton, and the mutant was screened and verified on the basis of the standard method (30). Likewise, the *S. candidus ΔprfE* mutant was constructed following the protocols described as above.

### *In vitro* assays of ForT and PrfE

Reactions consist of 50 mM Tris-Cl buffer (pH 8.0), 1.5 mM PRPP, 20 mM Mg^2+^ (or other related divalent metal ions), 1 mM compound **11/9a**, and 20 μg ForT/PrfE at 30 °C for 4 h. After that, reactions were terminated by the addition of an equivalent volume of methanol, and protein was removed by centrifugation. The HPLC analysis was performed on a reverse phase C18 column (Shimadzu, 5 μm, 4.6 × 250 mm) with an elution gradient of 10%-30% methanol: 0.1% aqueous TFA (HPLC grade) over 15 min at 0.5 mL/min, and then the methanol system returned to the initial 10% and maintained the ration for another 15 min. The elution was monitored at 290 nm (for ForT reaction) or 270 nm (for PrfE reaction) with a DAD detector, and the data were analyzed offline with Shimadzu data software.

## Accession number

The DNA sequence is available in the GenBank database under individual accession numbers KY705052, MH493900, and KY682079.

## ACKNOWLEDGEMENTS

We are grateful to Prof. Paul R. Jensen from UCSD (CA, USA) for kindly providing us with the strain *Salinispora arenicola* CNS-205. We also sincerely thank Prof. Pablo Sobrado from Virginia Tech (VA, USA) for the generous gift of the NbtG expression plasmid. This work was supported by grants National Key R & D Program of China (2018YFA0903203), and the National Natural Science Foundation of China (31770041).

